# Effects of latent toxoplasmosis on olfactory functions of men and women

**DOI:** 10.1101/231795

**Authors:** Jaroslav Flegr, Manfred Milinski, Šárka Kaňková, Martin Hůla, Jana Hlaváčová, Kateřina Sýkorová

## Abstract

The prevalence of toxoplasmosis is higher in schizophrenics than in the general population. It has been suggested that certain symptoms of schizophrenia, including changes in olfactory functions, are in fact symptoms of toxoplasmosis that can be easily detected in schizophrenics only due to the increased prevalence of toxoplasmosis in this population. Schizophrenics have impaired identification of odors and lower sensitivity of odor detection. Here we searched for differences in olfactory functions between 62 infected and 61 noninfected non-schizophrenic subjects. The infected men scored better in the standard odor-identification test. The infected women rated all smells as more intensive while the infected men rated nearly all smells as less intensive. Infected women rated the pleasantness of the smell of undiluted cat urine as higher than the non-infected women and the opposite was true for the men (the opposite direction shifts in men and women were described earlier for highly diluted cat urine). Toxoplasmosis had no effect on the rated pleasantness of the smell of other stimuli. Our results suggest that latent toxoplasmosis is associated with changes in the olfactory functions in humans; however, the observed changes differ from those observed in schizophrenics.

**Key findings:**

1. Infected men but not women show better odor identification ability than the non-infected controls.
2. The infected women rated all smells as more and men as less intensive than the controls.
3. The infected women rated smell of cat urine as more and men as less pleasurable than the controls.
4. Toxoplasmosis had no effect on the rated pleasantness of the smell of other stimuli.
5. We found no new evidence for the toxoplasmosis hypothesis of schizophrenia.

## INTRODUCTION

The protozoan parasite *Toxoplasma gondii* infects between 10-80% of the inhabitants of developed countries, depending on climate, hygienic and dietetic habits and the density of cats (the definitive host of *Toxoplasma*) in particular areas (Pappas *et al.*, 2009). After a short phase of acute toxoplasmosis, the disease proceeds into the life-long latent phase. For a long time, this phase was considered more or less asymptomatic in immunocompetent individuals. However, within the last 15 years, more and more results suggest that the latent form of toxoplasmosis is associated with increased frequencies of various disorders, mainly but not exclusively with neuropsychiatric disorders, for review see (Flegr *et al.*, 2014; Sutterland *et al.*, 2015). It is widely accepted now that toxoplasmosis plays an important role in the etiology of schizophrenia. The prevalence of toxoplasmosis is higher in schizophrenia patients (Torrey *et al.*, 2007; Torrey *et al.*, 2012), especially in those with a continuous course of the disorder (Celik *et al.*, 2015). *Toxoplasma* codes two enzymes for the syntheses of dopamine (Gaskell *et al.*, 2009), the neurotransmitter that plays a very important role in positive symptoms (hallucinations and delusions) of schizophrenia (Willner, 1997). Latent toxoplasmosis is associated with the increased concentration of this neurotransmitter in the brain of infected human (Flegr *et al.*, 2003) and rodent (Prandovszky *et al.*, 2011) hosts, which probably explains the higher intensity of positive symptoms and longer hospitalization in Toxoplasma-infected schizophrenics (Holub *et al.*, 2013; Wang *et al.*, 2006; Yolken *et al.*, 2009). The specific morphological changes of the brain of schizophrenia patients, namely the decrease of grey matter density bilaterally located within the caudate, median cingulate, thalamus and occipital cortex and in the left cerebellar hemispheres can be detected only in those patients who are infected with *Toxoplasma* (Horacek *et al.*, 2012).

It has been suggested that some characteristics of schizophrenia patients could in fact be the results of the *Toxoplasma* infection, which is more frequently documented in schizophrenia patients than in the general population, rather than the direct effect of schizophrenia (Priplatova *et al.*, 2014). It is known, for example, that both schizophrenia patients and people with latent toxoplasmosis have an increased latency of startle reflex and a decreased effect of prepulse on the latency in the acoustic startle reflex inhibition test (Pearce *et al.*, 2013; Priplatova *et al.*, 2014). It has been also described that both toxoplasmosis and schizophrenia are associated with characteristic changes of olfactory functions. Toxoplasma-infected subjects rate the pleasantness of smell of highly diluted urine of the definitive host of *Toxoplasma*, the cat, but not the pleasantness of smell of urine of other four species, differently than the non-infected controls (Flegr *et al.*, 2011). Namely, the *Toxoplasma* infected men rated this smell as more pleasant, which recalls the well-known “fatal attraction phenomenon” observed in infected rodents (Berdoy *et al.*, 2000; Vyas *et al.*, 2007a) and chimpanzee (Poirotte *et al.*, 2016), while the Toxoplasma-infected women rate the smell of the diluted cat urine as less pleasant. The change of the natural fear of the smell of cat predators of animals towards an attraction to this smell after the *Toxoplasma* infection is considered to be the product of manipulation activity of the *Toxoplasma* aimed to increase the chance of its transmission from the intermediate to definitive host by predation. It has been shown that epigenetic modification (demethylation) of regulatory parts of certain genes in the medial amygdala is probably responsible for the observed behavioral changes (Dass & Vyas, 2014). The schizophrenia-associated effects seem to be less specific (however, the “fatal attraction” effect has never been studied in this population). A strong negative influence of schizophrenia on the performance of patients in the odor identification test has been reported in many studies. A couple of studies also demonstrated schizophrenia-associated deficits in the detection threshold sensitivity, the odor discrimination and the olfactory recognition memory; for an excellent analytical review see (Moberg *et al.*, 1999). These olfactory functions have not been studied in people with latent *Toxoplasma* infection.

In the present study we searched for support for the hypothesis that the high prevalence of Toxoplasma-infected subjects (with olfactory function modified by the manipulation activity of the parasite) could be responsible for the reported olfactory deficits in schizophrenia patients. We compared the performance of Toxoplasma-infected and *Toxoplasma*-free men and women in two variants of the odor identification test and investigated the differences in rating the intensity and pleasantness of four odor samples of animal origin (zibet, moschus, ambra, cat urine) and one of plant origin (jasmine). *Toxoplasma* has mostly opposite effects on behavior and personality of men and women and often different effects on Rh positive and Rh negative subjects (Flegr, 2013). The behavioral and performance effects of toxoplasmosis also mostly increase with time since the infection. Therefore, our *Toxoplasma*-infected and *Toxoplasma-free* populations were matched for sex, age, and Rh.

## MATERIALS AND METHODS

### Subjects and procedure

Most of subjects were past students of biology who participated in various parasitological and evolutionary psychological studies that have run at the Faculty of Science in past 15 years. About one third were registered members of an internet community “Guinea pigs” (Pokusni kralici in Czech, www.facebook.com/pokusnikralici). This community of people willing to take part in diverse evolutionary psychological experiments consists of subjects of various ages, education levels, occupations, and locations (Flegr & Horáček, 2017). Within the last 4 years they were invited to come to the faculty or to other institutions in five large Czech cities for testing for toxoplasmosis and Rh phenotype. We invited by telephone the Toxoplasma-infected Rh negative and Rh positive subjects, as well as the same number of *Toxoplasma*-free controls balanced for sex, Rh, and age to come to the faculty to participate in a 45 minutes “sniffing experiment”. About 95% of subjects agreed to participate and about 75% were actually able to come to some of suggested appointments for testing. Men were tested within two weeks of January and two weeks of February 2017 and women within two weeks of March 2017. The participants were given the instructions not to use perfume, perfumed soap or perfumed deodorant on the day of the experiment and not to smoke or consume aromatic meals at least one hour before the session. The participants were not informed that the study was concerned with toxoplasmosis and this information was not written in the informed consent but was provided to the participants during a 5 minutes debriefing after the end of the experiment. After reading and signing the informed consent, the participants (alone and in their own tempo) rated the intensity and pleasantness of 10 odor samples in a quiet, ventilated room. One half of each subgroup rated the variant A (1 Jasmin, 2 Moschus, 3 Ambra, 4 Zibet, 5 only solvent, 6 Zibet, 7 Ambra, 8 Moschus, 9 Jasmin, 10 undiluted cat urine) and one half the variant B (1 Zibet, 2 Ambra, 3 Moschus, 4 Jasmin, 5 only solvent, 6 Jasmin, 7 Moschus, 8 Ambra, 9 Zibet, 10 undiluted cat urine). To rate the pleasantness of odors, the participants had to respond to the question “How pleasurable would it be to use perfume containing this ingredient and to smell like this?” using a graphic scale of the length 9 cm anchored with “very unpleasurable” on the left and “very pleasurable” on the right. To rate the intensity of odors, the participants had to respond to the question “How intensive is this smell” using a second graphic scale of the length 9 cm anchored with “very intensive” on the left and “very weak/I do not smell anything” on the right. To study the unconscious smell preference by the chromassociation method, the raters were given 12 color pencils and were asked to choose which color was best suited to a particular smell. After finishing this part of the test, the participants were asked to complete an anamnestic questionnaire lasting ten minutes and containing 24 questions mostly on health, moods, and using perfumes and deodorants. They were also asked how many cigarettes they usually smoke per day (occasional smokers were recommended to use decimal numbers). In the final part of the session, an assistant of the same sex presented them, one after one, 12 sniffing stick of the Burghart odor identification test. In the free-recalling variant of the test the assistant asked the participant what he thinks this smell is, recorded the answer and then in the standard variant of the sniffing test presented him a card with four alternative answers to be chosen from. After the end of the session, the raters finished the second part of the chromassociation test, i.e., they arranged the pencils from the most favorite to the least favorite color. Then the assistant checked the completeness of the material and explained aims of the study to the participant. Each rater was invited separately for a particular time and got no reward for his participation, except a special badge issued for this occasion and sometimes the compensation of travel expenses for the non-Prague participants. The project, including the method of subjects' recruitment, content and form of informed consent, and procedure of the data handling, has been approved by the Institutional Review Board of the Faculty of Science, Charles University (Etická komise pro práci s lidmi a s lidskym materiálem Přírodovedecké fakulty Univerzity Karlovy), approval number 2016/27.

### Material

The attractiveness of cat urine for *Toxoplasma*-infected and *Toxoplasma*-free subjects (and rodents and chimpanzees) is known to differ. In our recent historical past, nearly the whole human population was infected with *Toxoplasma*. It could therefore be expected that the attractiveness of some perfume ingredients of animal origin could also be the result of the *Toxoplasma* manipulation activities – of reprogramming of specific neural circuits in the amygdala. Therefore, we used three perfume ingredients of animal origin (zibet, moschus, ambra) and one of plant origin as a control (jasmine).

### Perfume ingredients

The fragrances were natural products of the highest quality.

#### Zibet

1g Zibet (Zibet absolue Essencia 60-4250-0) in 80ml base mixing (70g Ethanol 99.8% + 10g Aqua dest.) pestled in Achat mortar, transferred to a Duran flask, 1 drop Tween 80 added, for 24h in shaking water quench (50°C/120rpm), then filtered (5-8μm). This preparation was diluted further 1:2.5: 1.6ml Zibet (1:65 as described above) + 2.4ml base mixing. The final dilution for the experiment was 1:162.5.

#### Ambra

1g Ambra (Ambra grau Essencia 60-2350-0) in 80ml base mixing, pestled in Achat mortar, transferred to a Duran flask, 1 drop Tween 80 (P4780 Sigma) added, for 24h in shaking water quench (50°C/120rpm), then filtered (5-8μm). This base preparation was not diluted further; the final dilution for the experiment was 1:65.

#### Moschus

0.8ml Original product (Moschuskörner 15% in Ethanol, Primavera 11128) + 3.2ml base mixing. The final dilution for the experiment was 1:5.

#### Jasmin

20μl Original product (Jasmin absolue, Primavera 10150) + 6.38ml base mixing. The final dilution for the experiment was 1:320.

Number-coded glass vials each of which contained a 3 cm long smelling strip (as used in perfumology) that was fixed to the plug. Each strip was supplied with two drops (about 27 μl) of the respective perfume ingredient.

#### Cat urine

Aliquots of the mixture of urine from five female cats were frozen at -18° C. One week before the experiment, 10 μl of the mixture for women and 20 μl for men were pipetted on the strip of filtrating papers in the testing vial and the vials were again put into a -18°C freezer. The larger volume of urine was used in men because two of six men did not identify the presence of any odor in our pilot experiment. At 10 minutes before the experiment, the vials were taken from the freezer and kept at room temperature until the beginning of the test.

### Odor identification tests

The Burghart Sniffin’ Sticks Screening 12 Test (Medisense, The Netherlands) with peppermint, fish, coffee, banana, orange, rose, lemon, pineapple, cinnamon, cloves, leather and liquorice odors was used in two variants of the odor identification test. Each participant was at first handed one sniffing stick by an assistant of the same sex (in the same sequence from 1 to 12) to try to identify a particular odor (the free-recalling variant of the test). Then, in the standard variant of the test, the assistant handed him/her a multiple choice form containing names of four different odors and the participant tried to identify the correct one. The assistant recorded the answers of the participants in both parts of the odor identification test and handed the participants the next sniffing stick.

### Immunological tests for T. gondii infection and Rh phenotype

All testing was performed at the National Reference Laboratory for Toxoplasmosis, National Institute of Public Health, Prague. The complement-fixation test (CFT), which determines the overall levels of IgM and IgG antibodies of particular specificity, and Enzyme-Linked Immunosorbent Assays (ELISA) (IgG ELISA: SEVAC, Prague) were used to detect the *T. gondii* infection status of the subjects. ELISA assay cut-point values were established using positive and negative standards according to the manufacturer's instructions. In CFT, the titer of antibodies against *T. gondii* in sera was measured in dilutions between 1:8 and 1:1024. The subjects with CFT titers between 1:8 and 1:128 were considered *T. gondii* infected. Only subjects with clearly definitive results of CFT or IgG ELISA tests were diagnosed as *T. gondii*-infected or *T. gondii*-free. A standard agglutination method was used for Rh examination. A constant amount of anti-D serum (monoclonal human IgM blood grouping reagents; antiD clones MS-201, RUM-1, Millipore (UK) Ltd.) was added to a drop of blood on a white glass plate. Red cells of Rh-positive subjects were agglutinated within 2–5 minutes.

### Statistics

Before the analyses, the ratings on a graphical scale were measured and recorded independently by two persons and any inconsistencies were checked and corrected. They also calculated unconscious smell preference scores by computing the mean attractiveness score of color attributed by a particular rater to individual odors – the color the most attractive for the rater was coded as 12, the most unattractive as 1. To control for effects of sequence of presentation each perfume ingredient was present in two vials, therefore the arithmetic mean was calculated from these two values. Statistica v. 10.0 was used for all tests t-tests, Pearson’s and Spearman correlation test, ANCOVA, repeated measure ANCOVA, and for testing of the tests’ assumptions. Partial Kendall correlation with age as the confounding variable was computed using an Excel sheet available at http://web.natur.cuni.cz/flegr/programy.php. Some variables, e.g. the frequencies of smoking, had highly asymmetric distribution, therefore we used both parametric and nonparametric tests for their analysis; however, the results were qualitatively the same. The Bonferroni’s correction was used for the correction for multiple tests. The data file is available at https://figshare.com/s/f112b4805f6e7c059819.

## RESULTS

The final sample consisted of 60 women (age: 28.0, SD 4.46) and 63 men (age: 28.1, SD 5.69). In the women, 30 subjects were *Toxoplasma*-infected (15 Rh-negative and 15 Rh-positive) and 30 *Toxoplasma*-free (15 Rh-negative and 15 Rh-positive). In the 63 men, 32 subjects were *Toxoplasma-*infected (9 Rh-negative and 23 Rh-positive) and 31 were *Toxoplasma*-free (8 Rh-negative and 23 Rh-positive). No significant differences in age existed between the men and women, *Toxoplasma*-infected and *Toxoplasma*-free, Rh-negative or Rh-positive groups of subjects (all t-test p values > 0.20).

### Effect of toxoplasmosis on the identification of odors

Women recognized more odors than men in both variants of the test (p = 0.01, η^2^ = 0.090). The Toxoplasma-infected subjects identified more odors in the standard variant of the test (selection of the answer from four different options) and fewer odors in the first free-recalling variant of the test (Fig. 1). The effect of the interaction toxoplasmosis-type of the test was significant when the variable Rh was present (p = 0.030, η^2^ = 0.041) or absent (p = 0.016, η^2^ = 0.048) in the model. A separate analysis of the two variants of the test showed that the effect of toxoplasmosis was significant in the standard variant of the test (Rh included: p = 0.048, η^2^ = 0.034); Rh not included: p = 0.034, η^2^ = 0.037) and not significant in the free-recalling variant of the test (Rh included: p = 0.473, η^2^ = 0.005; Rh not included: p = 0.406), η^2^ = 0.006). Partial Kendall correlation with the age as a confounding variable showed that the positive effect of toxoplasmosis in the standard variant of the test was significant for all participants (partial Tau = 0.140, p = 0.021) and men (partial Tau = 0.242, p = 0.005), but non significant for women (partial Tau = 0.078, p = 0.379).

**Fig. 1.**
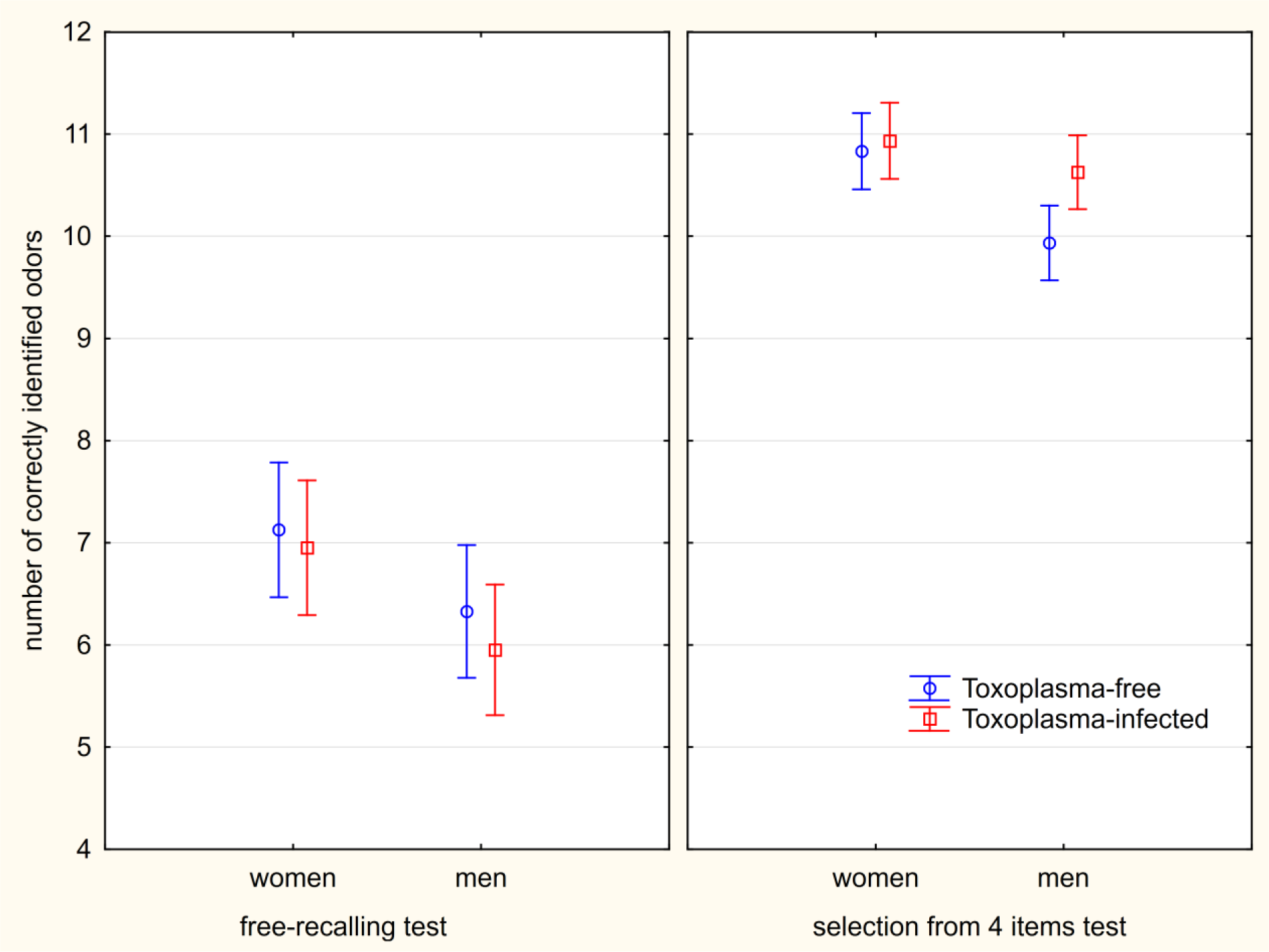
Differences of abilities to recognize odors in the *Toxoplasma*-free and Toxoplasma-infected subjects. *The error bars show 95% confidence intervals.*

### Effect of toxoplasmosis on rated intensity of odor

A negative correlation existed between the rated intensity and pleasantness of particular smells as well as between the age or frequency of smoking and the rated intensity of smell. Also, positive correlation existed between frequency of smoking and rated pleasantness of smell, see Table 1. Therefore, we included the covariates of the age of subjects and the frequency of smoking into all analyses, and the age of subjects, frequency of smoking and intensity of smell rated by the same subject into all analyses in which the effects of toxoplasmosis on pleasantness of smell was studied.

**Table 1.** Correlation between rated pleasantness and intensity of particular odors

|  | age | Cat<br>intensity | Jasmin<br>intensity | Moschus<br>intensity | Ambra<br>intensity | Zibet<br>intensity | neutral<br>intensity | smoking<br>frequency |
| --- | --- | --- | --- | --- | --- | --- | --- | --- |
| age | 1.00 | -0.01 | -0.13 | -0.10 | -0.11 | **-0.23** | **-0.31** | -0.11 |
| Cat | 0.02 | **-0.67** | **-0.21** | **-0.18** | -0.04 | -0.05 | -0.15 | 0.11 |
| Jasmin | **-0.19** | **-0.21** | **-0.28** | -0.10 | -0.13 | -0.05 | **-0.21** | 0.15 |
| Moschus | 0.00 | -0.06 | -0.12 | **-0.41** | -0.07 | -0.12 | -0.15 | 0.03 |
| Ambra | -0.05 | -0.05 | -0.16 | -0.10 | **-0.53** | **-0.42** | -0.12 | **0.19** |
| Zibet | 0.04 | 0.00 | **-0.22** | **-0.24** | **-0.47** | **-0.65** | -0.16 | 0.11 |
| neutral | -0.02 | -0.10 | -0.08 | -0.07 | -0.17 | **-0.18** | **-0.29** | 0.15 |
| smoking | -0.11 | -0.06 | 0.07 | 0.08 | **-0.24** | -0.10 | -0.02 | 1.00 |

Table shows Spearman *R* for the correlations of intensities (columns) and pleasantness (rows) of particular odors (and age and frequency of smoking). Significant coefficients ( $p < 0.05$ ) are printed in bold.

Repeated measure GLM with attributed intensity of 6 odors (including the neutral sample – the empty vial) as repeated measures and toxoplasmosis, sex, Rh, age, and intensity of smoking as the independent variables revealed the significant effect of toxoplasmosis-sex interaction (p = 0.015, η^2^ = 0.051), age (p = 0.005, η^2^ = 0.069), sex (p = 0.012, η^2^ = 0.055), age-type of odor interaction (p = 0.001, η^2^ = 0.035), sex-type of odor interaction (p < 0.0005, η^2^ = 0.135), smoking-type of odor interaction (p = 0.006, η^2^ = 0.028), and Rh-type of odor interaction (p = 0.017, η^2^ = 0.024). The Toxoplasma-infected women rated the intensity of the smell of all odors as higher (cat: partial Tau = 0.176, p = 0.047; jasmin: partial Tau = 0.060, p = 0.501; moschus: partial Tau = 0.179, p = 0.043; ambra: partial Tau = 0.164, p = 0.063; zibet: partial Tau = 0.191, p = 0.031; neutral: partial Tau = 0.180, p = 0.042), while *Toxoplasma*-infected men rated all except one of the intensities as lower (cat: partial Tau = -0.066, p = 0.444; jasmin: partial Tau = -0.081, p = 0.350; moschus: partial Tau = - 0.096, p = 0.269; ambra: partial Tau = 0.052, p = 0.547; zibet: partial Tau = -0.086, p = 0.318; neutral: partial Tau = -0.053, p = 0.539), see the Fig. 2.

**Fig. 2.**
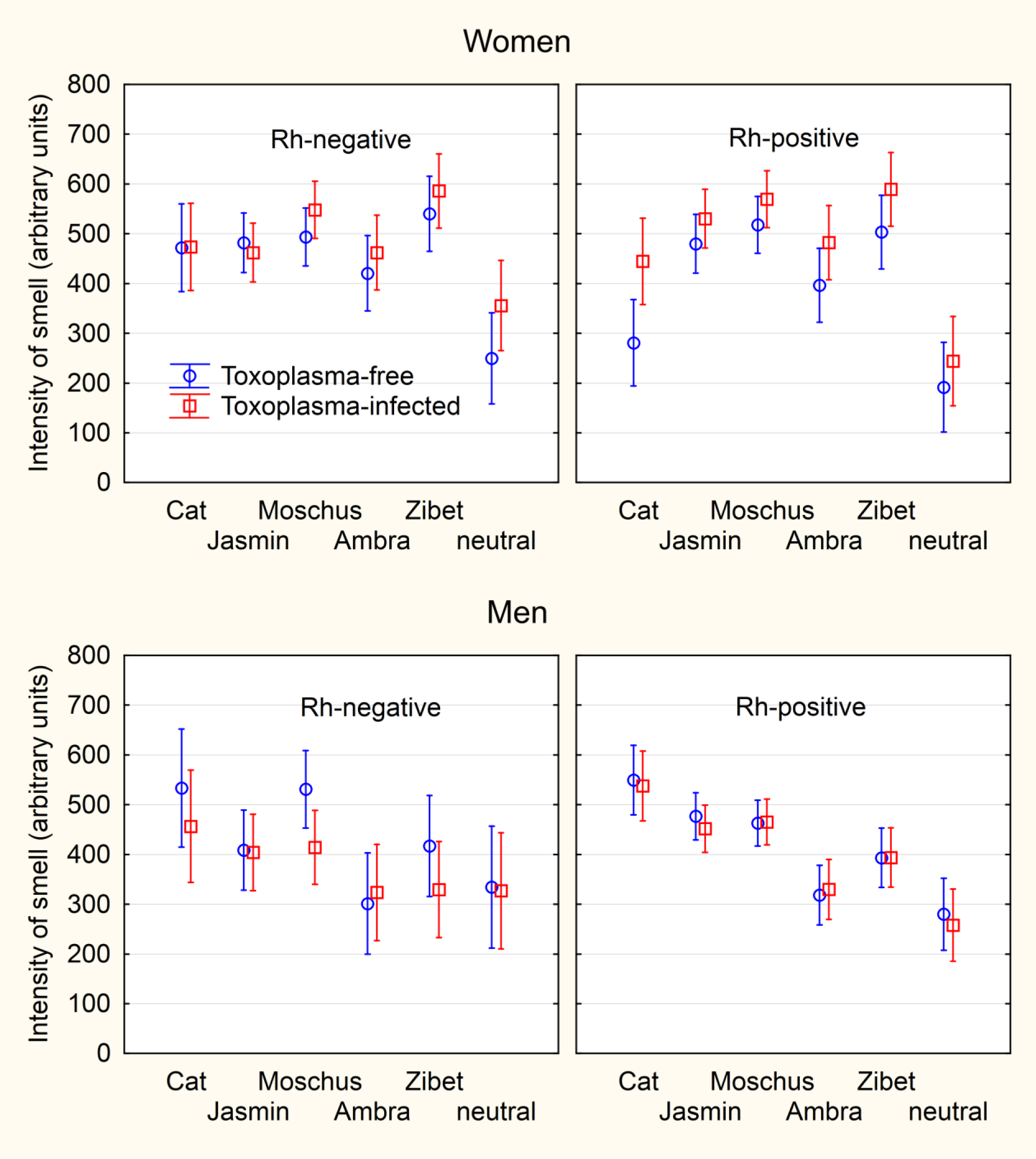
Difference in the intensities of smells attributed by women and men to particular odors. *The error bars show 95% confidence intervals.*

### Effects of toxoplasmosis on rated pleasantness of odors

Repeated measure GLM with attributed pleasantness of six odors (including the neutral sample) as repeated measures and toxoplasmosis, sex, Rh, intensity of smoking and age as an independent variables revealed only significant effects of sex (p = 0.009, η^2^ = 0.059), type of odor (p < 0.0005, η^2^ = 0.056), sex-type of odor interaction (p < 0.0005, η^2^ = 0.060), and a nearly significant effect of smoking (p = 0.056, η^2^ = 0.032), Fig. 3. No other effect was significant and the same was true for the simpler model without Rh. The results of separate analyses for particular odors with toxoplasmosis, sex, age and the intensity of particular smell as the independent variables provided a significant effect of toxoplasmosis-sex interaction on the smell of cat urine (p = 0.020, η^2^ = 0.046). The models that also included Rh and its interactions with other independent binary variables showed significant effects of toxoplasmosis-sex interaction for the cat urine (p = 0.015, η^2^ = 0.051), toxoplasmosis-Rh interaction for the moschus (p = 0.015, η^2^ = 0.051), and toxoplasmosis-sex-Rh interaction for the ambra (p = 0.030, η^2^ = 0.041) and the zibet (p = 0.036, η^2^ = 0.039). After the correction for multiple tests, however, no main effect of toxoplasmosis or of interaction with toxoplasmosis was significant in any model.

Identical analyses have been repeated with the pleasantness of odors estimated with chromatic association method. Except the effect of the type of odor (p = 0.001, η^2^ = 0.027), no other effects were significant.

**Fig. 3.**
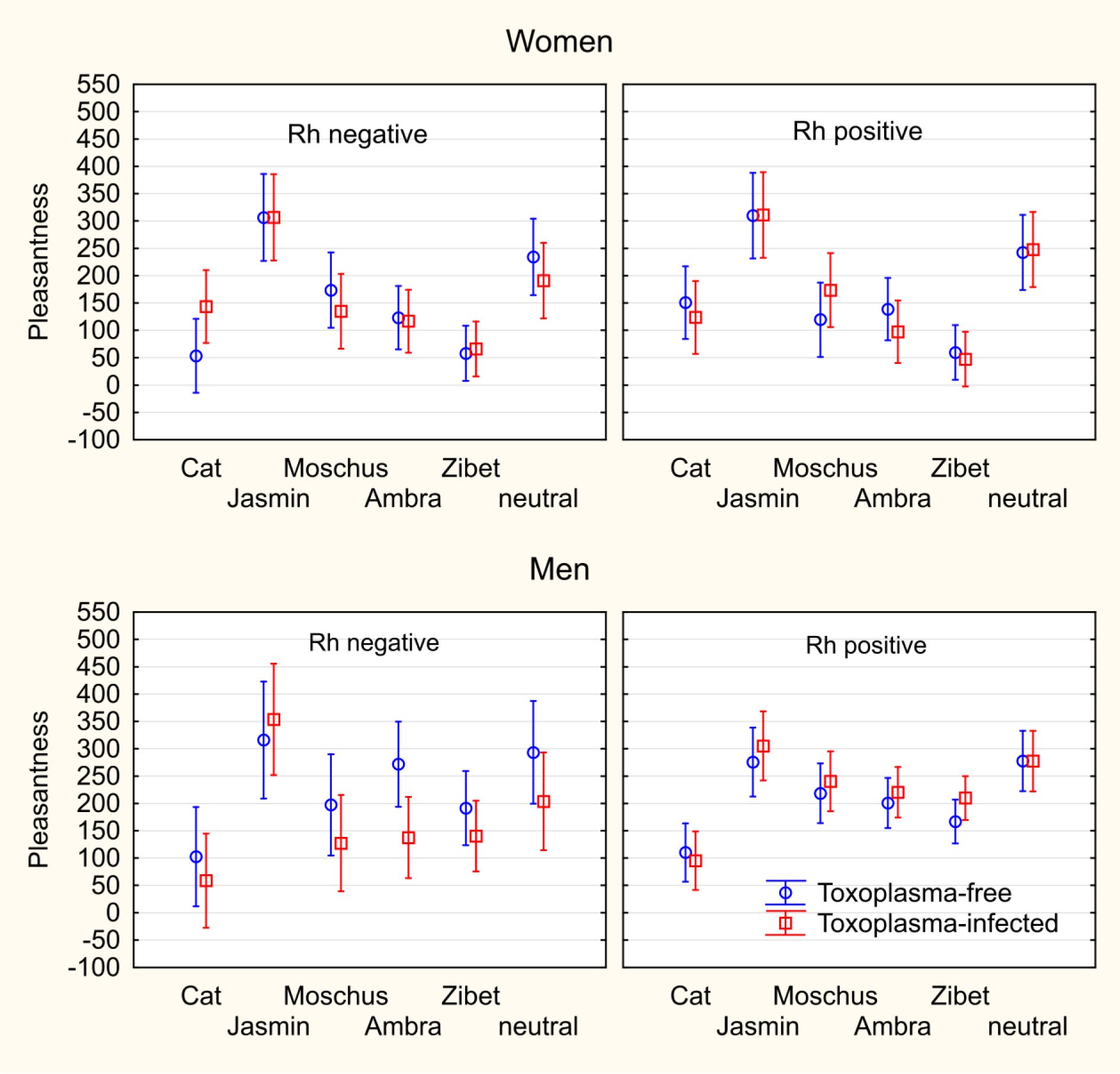
Difference in pleasantness attributed to particular odors by *Toxoplasma*-free and *Toxoplasma*-infected raters. *The figure shows mean pleasantness of odors in arbitrary units measured as the responses of the raters to the question “How pleasurable would it be to use perfume containing this ingredient and to smell like this?” (graphic scale, 9 cm). The spreads shows 95% confidence interval.*

**Fig 4.**
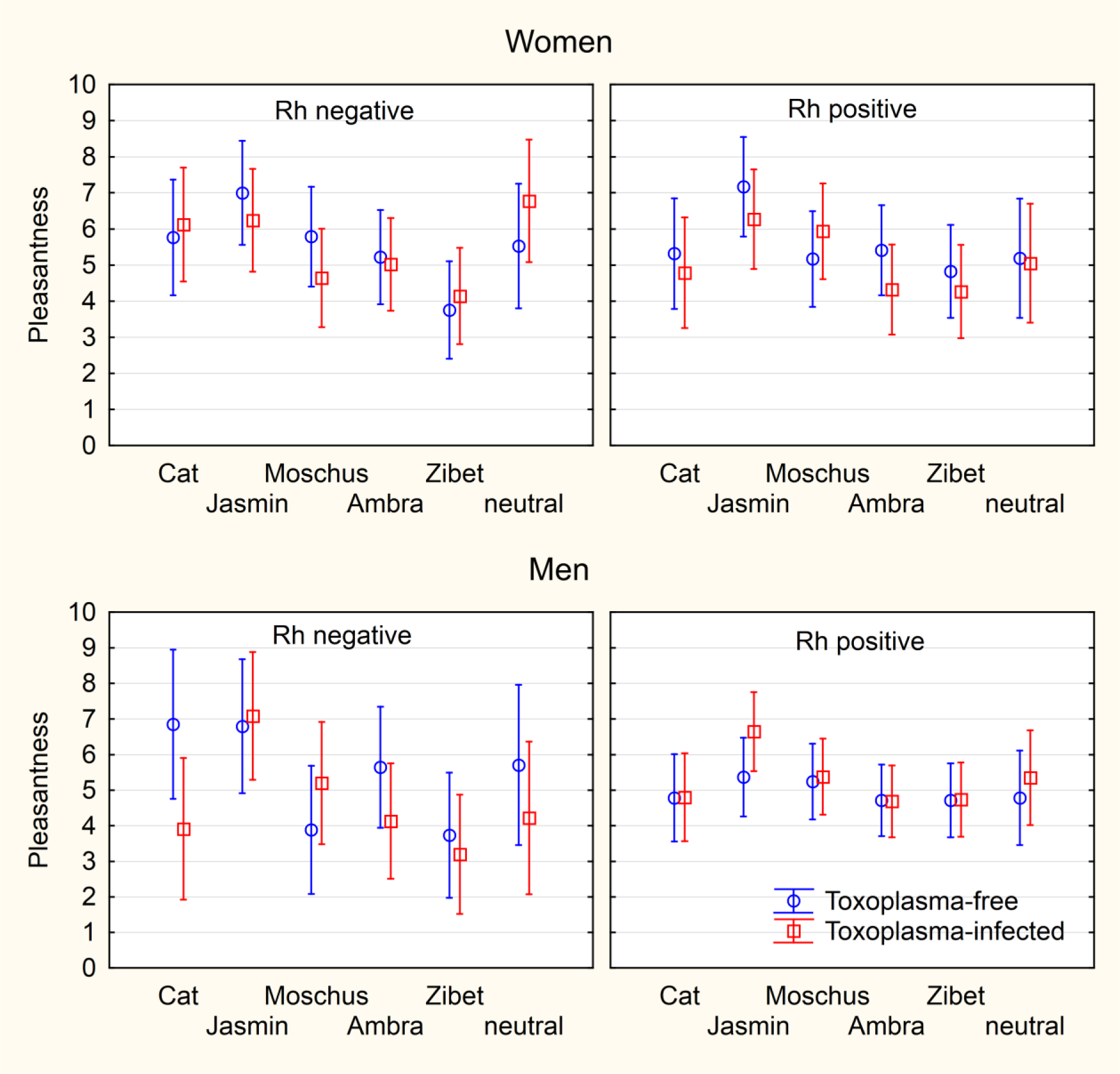
Pleasantness of odors estimated by the chromassociation method. *The figure shows mean pleasantness of color (scale 1-12) attributed by the individual raters to particular odor. The spreads shows 95% confidence interval.*

## DISCUSSION

We found significantly better performance of *Toxoplasma*-infected subjects, especially the men, in the standard odor identification test. *Toxoplasma*-infected women rated all smells as more intensive while the *Toxoplasma*-infected men rated nearly all smells as less intensive. Infected female raters rated the pleasantness of the smell of the cat urine as higher than the non-infected female raters and the opposite was true for the male raters. We found no evidence for the effect of toxoplasmosis on the rated pleasantness of the smell of jasmin, moschus, ambra, and zibet.

The better performance of the *Toxoplasma*-infected subjects in the odor identification test and the higher scores of intensity of smell attributed to the odors in women was unexpected. It is, however, possible that the infected subjects have in fact unchanged olfactory functions but try more hardly to identify the odor during the test and express higher willingness to rate the intensity of smell as higher. For example, infected women are known to express higher affectothymia, warmth, and cooperativeness than non-infected women (Flegr *et al.*, 1996; Lindová *et al.*, 2010). All these traits could results in larger effort invested into the test and in higher ratings attributed to particular odors. Infected men express higher concentration of testosterone (Flegr *et al.*, 2008a) and therefore higher competitiveness, which could also result in higher number of correctly identified odors, and higher suspiciousness (Lindová *et al.*, 2006), which could result in their lower ratings of the intensity of odors – they could suspect that some vials with low-intensity smell samples are empty).

The opposite effect of toxoplasmosis on the rated pleasantness of cat urine in men and women has been already described. However, in the previous study the infection increased the rated pleasantness of the smell in men and decreased it in women (Flegr *et al.*, 2011). The main difference between these two studies was that highly diluted urine was used in the first study and undiluted urine in the present study. It is known that the relation between the attractiveness of the smell of the cat urine for the *Toxoplasma*-infected rodents has a reverse U shape (Vyas *et al.*, 2007b). It is high for medium concentration of urine but low for very low and very high concentrations of urine. It will be necessary to include several concentrations of the cat urine into future studies and examine the effects of sex-toxoplasmosis-concentration interaction.

Formally, we did not detect any effect of toxoplasmosis on the rating of the pleasantness of odors of four perfume ingredients as all effects disappeared after the correction for multiple tests. There were just minor differences between the patterns when the pleasantness of odors was rated by the subjects or measured with the chromassociation test, except relative higher scores for cat urine obtained with the chromassociation tests. Based on results obtained with our set of wide spread natural ingredients we conclude that toxoplasmosis has no significant effect on the preference for these perfume ingredients to be used for themselves. The preference for these ingredients has been shown to correlate with the possession of specific MHC immune genes (Milinski & Wedekind, 2001) and MHC genes play a role in resistance of intermediate hosts to toxoplasmosis (Fux *et al.*, 2003; Mack *et al.*, 1999). The absence of significant effects of toxoplasmosis on the pleasantness of odors suggests that different MHC genes play a role in the preference of perfumes and in the resistance to *Toxoplasma*.

The negative results of this part of the study, namely the absence of any Rh-toxoplasmosis interaction, can be, at least partly, caused by a relatively low number of participants of the study, especially the low number of Rh-negative men (8 *Toxoplasma*-free and 9 *Toxoplasma*-infected). Within past 25 years, we have tested many thousands of volunteers for toxoplasmosis and about three thousands of them are willing to continue to participate in our experiments. However, the number of men is lower than number of women due to unbalanced sex ratio in local biology students. Further, the prevalence of toxoplasmosis in current students decreased to less than 10 % and the frequency of Rh-negative persons in the Czech population is about 16 %. Therefore, we were not able to convince more *Toxoplasma*-infected, Rh negative men to come to our lab for testing.

Rh factor has a very strong effect on the response of human organisms to *Toxoplasma* infection. Some effects of toxoplasmosis are much stronger in Rh negative subjects and some can be observed only in this population or in Rh-positive homozygotes (Flegr *et al.*, 2009; Flegr *et al.*, 2008b; Kaňková *et al.*, 2010; Novotná *et al.*, 2008). In the present study we did not detect any interaction of Rh-toxoplasmosis that would survive the correction for multiple tests. However, we found that the Rh negative and Rh positive subjects differ in the intensity and possibly also pleasantness of odor attributed to different perfume ingredients, see Fig. 2. This anecdotal observation could have some practical implication for the perfume industry and would deserve future attention.

The main topic of the present study was the effect of toxoplasmosis on the olfactory function of healthy men and women. However, we found some unrelated known or unknown phenomena that deserve future attention. Our data suggest that smoking has positive influence on the on rating pleasantness of all odors and negative influence on rating intensity of some, but not all odors. The subjectively rated intensity to, as well as the pleasantness of, some odors differs between men and women. The women rated the pleasantness of the smell of moschus, ambra and especially zibet very low not only in comparison with the smell of a neutral sample but even in comparison with the smell of the cat urine. During the rating sessions, the participants were requested to answer the question “How pleasurable would it be to use a perfume containing this ingredient and to smell like this?” While men strongly disliked the prospect of smelling like cat urine, the women rated the smell of cat urine as more pleasant than that of zibet, the smell of which is subjectively reminiscent of cat urine. This seems to be in conflict with the observation showing that zibet is contained in 31%, i.e. 152 of 489, of perfumes for women but only in 6 %, i.e. 23 of 377, of perfumes for men (data analyzed from H&R Fragrance Guide - Fragrances on the international Market. Glöss, Hamburg 1995). The most obvious explanation, namely that the perfumes for men are mainly chosen by women and vice versa, is almost certainly false – perfumologists regard it as a fact that both women and men choose perfumes for themselves and usually do not use any other perfumes chosen by someone else (e.g., Perfumery – The Psychology and Biology of fragrance. Steve Van Toller, George H. Dodd (eds). Chapman & Hall, London, 1994).

### Limitations

The most important limitation is a rather low number of Rh negative men participating in the study. Due to this limitation, all positive and negative results concerning the effects of and the interaction with Rh factor must be considered only preliminary. We used only the small variant of the sniffing test. Many women identified all 12 samples in the standard variant of the test. Therefore, the sensitivity of the method was decreased due to ceiling effect in women and also partly in men. We used only one concentration (and one sample of) the cat urine in the set of samples. In future studies, both undiluted and highly diluted samples of cat urine as well as a sample of urine of an animal that is not the definitive hosts of *Toxoplasma gondii* should be used.

## Conclusions

Our data confirmed that latent toxoplasmosis has some effects on the olfactory function of humans. The strength of the effects is mostly low (η^2^ < 0.06), which corresponded to strength of other effects of toxoplasmosis on human performance and behavior. In contrast to other toxoplasmosis-associated effects, no significant differences were observed between Rh negative and Rh positive infected subjects. In contrast, an opposite effect of toxoplasmosis on the olfactory functions was observed in men and women, the phenomenon which has been described earlier for many behavioral and personality traits (Flegr *et al.*, 1996; Lindová *et al.*, 2006). This suggests that the observed differences, possibly including the observed analogy of the fatal attraction phenomenon (Flegr *et al.*, 2011), are not the primary effects of toxoplasmosis on the olfactory system but rather the secondary effects of toxoplasmosis-associated personality changes. According to the stress coping hypothesis (Lindová *et al.*, 2010), the opposite effects of toxoplasmosis on the behavior and personality of men and women can be explained by the opposite behavioral reaction of men and women to the chronic stress that is caused by the life-long parasitic infection. It is known that stressed men use more individualistic and antisocial (e.g. aggressive, hostile) forms of coping with stress, while stressed women are more likely to seek and provide social support. The observed difference in performance of infected and non-infected men and women could be just secondary effects of such coping.

We detected some differences in the olfactory functions of infected and non-infected subjects, however, the observed changes seems to differ from the changes described in schizophrenia patients. While schizophrenia patients have an impaired ability to identify odors, the *Toxoplasma* infected subjects performed better in the standard odor identification test. Therefore, our results are in conflict with our main hypothesis suggesting that the observed changes in olfactory performance of schizophrenia patients could be an experimental artifact caused by higher prevalence of *Toxoplasma*-infected subjects in the schizophrenia patients than in the non-clinical population (Priplatova *et al.*, 2014).

## ACKNOWLEDGMENTS

We thank Charlie Lotterman for the final revisions of our text.

